# AbDiver – A tool to explore the natural antibody landscape to aid therapeutic design

**DOI:** 10.1101/2021.11.03.467080

**Authors:** Jakub Młokosiewicz, Piotr Deszyński, Wiktoria Wilman, Igor Jaszczyszyn, Rajkumar Ganesan, Aleksandr Kovaltsuk, Jinwoo Leem, Jacob Galson, Konrad Krawczyk

## Abstract

**Motivation:** Rational design of therapeutic antibodies can be improved by harnessing the natural sequence diversity of these molecules. Our understanding of the diversity of antibodies has recently been greatly facilitated through the deposition of hundreds of millions of human antibody sequences in next-generation sequencing (NGS) repositories. Contrasting a query therapeutic antibody sequence to naturally observed diversity in similar antibody sequences from NGS can provide a mutational road-map for antibody engineers designing biotherapeutics. Because of the sheer scale of the antibody NGS datasets, performing queries across them is computationally challenging.

**Results:** To facilitate harnessing antibody NGS data, we developed AbDiver (http://naturalantibody.com/abdiver), a free portal allowing users to compare their query sequences to those observed in the natural repertoires. AbDiver offers three antibody-specific use-cases: 1) compare a query antibody to positional variability statistics precomputed from multiple independent studies 2) retrieve close full variable sequence matches to a query antibody and 3) retrieve CDR3 or clonotype matches to a query antibody. We applied our system to a set of 742 therapeutic antibodies, demonstrating that for each use-case our system can retrieve relevant results for most sequences. AbDiver facilitates the navigation of vast antibody mutation space for the purpose of rational therapeutic antibody design and engineering.

**Availability:** AbDiver is freely accessible at http://naturalantibody.com/abdiver.

## 1 Introduction

Monoclonal antibodies are the largest class of biotherapeutics. Development of successful antibody therapeutics requires selection and engineering of candidate sequences with favorable functional and developability properties. Knowledge of biologically possible mutations at specific positions can be employed to engineer biophysical properties of these molecules (Venkataramani *et al*., 2020). Next-generation sequencing (NGS) now allows us to capture millions of naturally-sourced B cell receptor (BCR) sequences in a single experiment, providing insight into natural antibody diversity (Kovaltsuk *et al*., 2018).

The richness of NGS data has implications for rational selection and design; models trained on these data have already shown promise for humanization (Marks *et al*., 2021) and binding prediction (Mason *et al*., 2021). It has also been shown that close sequence matches to clinically approved antibodies can be found in NGS datasets (Krawczyk *et al*., 2019), and clinically approved antibodies contain engineered mutations that can be recapitulated using the natural diversity from these datasets (Petersen *et al*., 2021). Exploration of natural antibody diversity from NGS datasets relative to a candidate therapeutic would therefore facilitate rapid and effective antibody engineering.

The volume of the publicly available antibody NGS data, makes investigation of their diversity mostly constrained to time-consuming bioinformatic endeavors. Available solutions either focus solely on clonotypes (Jones *et al*., 2021), CDR3s (Zhang *et al*., 2020) or do not allow for fuzzy matches (Olsen *et al*., 2021). To tackle such issues, we created AbDiver, an online tool allowing users to discover and characterize the natural sequence diversity surrounding a query antibody of interest. We provide three different approaches for performing this mapping 1) annotation of the natural positional diversity statistics on a position by position basis for a given query sequence 2) finding close matches to a full variable region sequence and 3) identifying sequences that would be classed as belonging to the same clonotype as the query sequence (based on CDR3 and germline V/J gene segments). AbDiver thus provides an accessible way to find a natural reference for a query therapeutic sequence, offering insights for sequence selection and rational design.

## 2 Implementation & benchmarking

### Data

As the underlying data we used publicly curated unpaired BCR NGS datasets from the Observed Antibody Space (OAS) (Kovaltsuk *et al*., 2018). In May 2021, the dataset encompassed 81 studies with 906,933,358 (105,730,531 light chains and 801,202,827 heavy chains) unique BCR sequences numbered according to the IMGT scheme (Lefranc *et al*., 2003). We envisage updates to the services as more datasets become available. For benchmarking, we employed a set of 742 therapeutic antibodies, discontinued, in clinical trials or approved in USA and the EU extending a set from our previous study (Krawczyk *et al*., 2021). Certain therapeutics were multi-specific or contained chain duplicates resulting in 738 unique heavy chains, 707 unique light chains, 686 unique CDRH3s and 573 unique CDRL3s.

### V-region profiling service

The AbDiver V-region natural profiling service annotates a query variable region antibody sequence with the naturally observed amino acid frequency statistics for each position (Figure 1). The frequency statistics are calculated from all antibodies comprising the same combination of V-gene and J-gene. Separate statistics were calculated for genes and their constituent alleles to allow for fine-grained allelic analysis, but also to reflect the ongoing effort in allele annotation (Smakaj *et al*., 2020). Each IMGT position in each profile contains statistics from amino acid frequencies calculated for each study separately. Amino acid positional frequency for a study was incorporated if it included at least 100 observations at a given position. For each position, we calculated the study-specific Shannon entropy and ranks of the amino acids by frequency. Query sequence sharing the germline genes/alleles of a given profile is then annotated at each IMGT position with the ranks and entropies of the given amino acid, averaged from the ranks and entropies of individual studies. This approach is designed to mitigate the effects of different numbers of sequences, techniques and disease states contributed by different studies, emphasizing frequency commonalities independent of study-specific biases.

**Figure 1.**
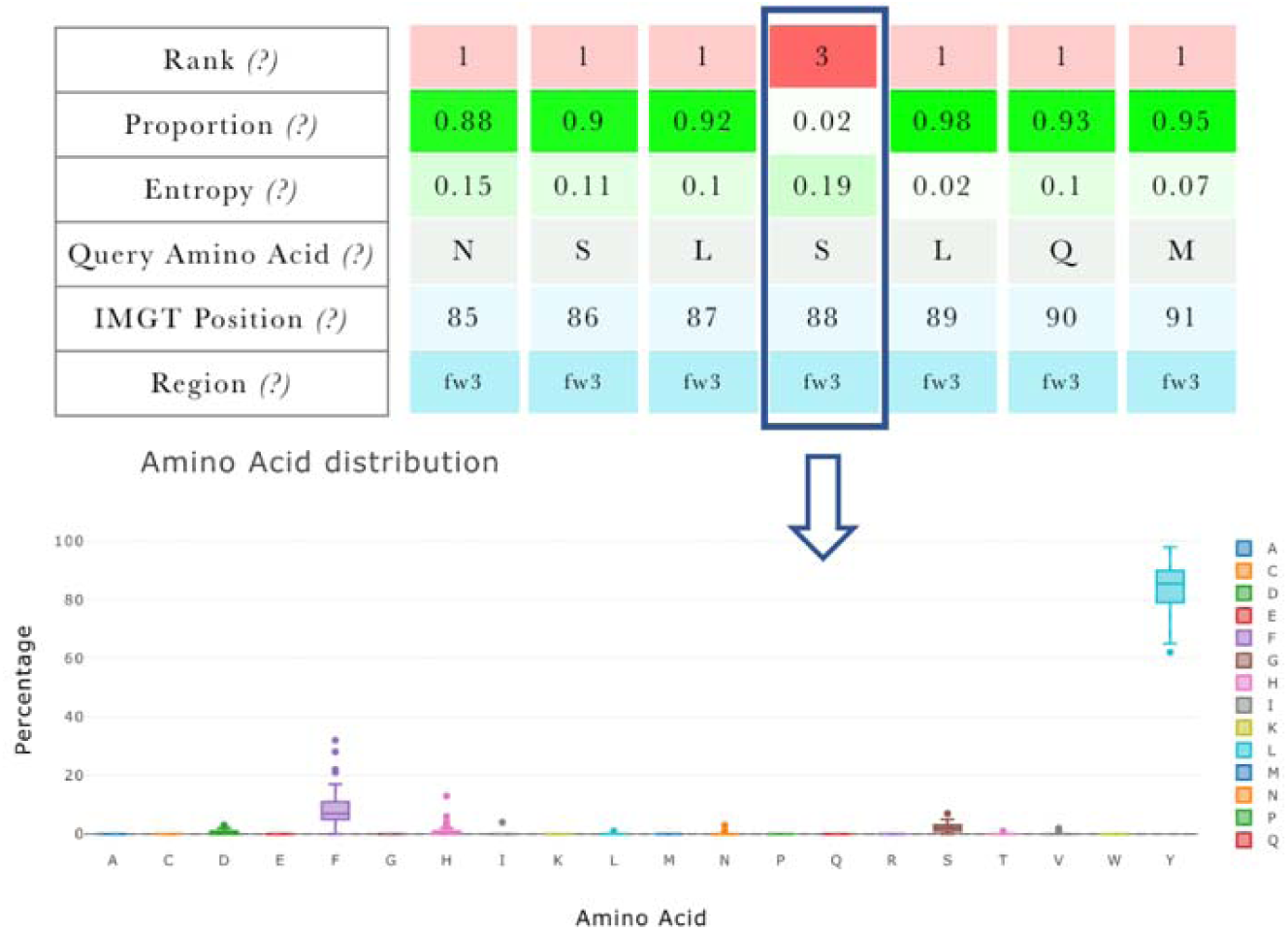
Visualization of the profiling service. Query sequence is compared to the frequency distribution statistics of amino acids of NGS sequences within the same gene or allele. Statistics reflect frequency commonalities drawn from independent studies.

### V-region profile benchmark

We tested whether the profiling service could find suitable profiles for our 742 therapeutics. A successful profile was arbitrarily defined as requiring at least 10,000 sequences contributing to its calculation (supplementary Figures 1, 2). Using allele-based profiles there were 688/738 (93.22%) heavy chains and 486/707 (68.74%) light chains where profiles had more than 10,000 OAS sequences. Using the less-stringent gene-based profiles there were 699/738 (94.71%) heavy chains and 496/707 (70.15%) light chains where profiles had more than 10,000 sequences. The smaller number of light chains we can find suitable profiles for results from a skew towards heavy chains in NGS depositions. Our profiles indicated that the majority of therapeutic antibodies contain framework mutations that are not commonly found in naturally occurring antibodies. In total, 191/738 (25.88%) heavy chains and 125/707 (17.68%) light chains from the therapeutic antibodies contained more than five framework mutations not commonly found in naturally occurring antibodies (supplementary Figures 3, 4). Our service identifies profiles for the majority of therapeutic antibodies highlighting non-trivial positional frequency information.

### Sequence retrieval service

We created k-mer (k=5) based indexes for CDRs in full variable-region sequences and CDR3s separately. Variable sequence matches are identified based on the same length CDR1, CDR2 with one residue discrepancy allowed for CDR3. Clonotypes are identified on the basis of the same V-gene and CDR3 sequence identity. The average retrieval time for heavy variable sequences is 4.40s and for heavy CDR3s 2.68s (see supplementary Figure 5). The matches are IMGT aligned and presented using Multiple Sequence Alignment (Martin, 2014). Results of both searches are presented using interactive tables highlighting the leading themes in the studies (e.g., studied disease, vaccine) facilitating further exploration of results.

### Sequence retrieval benchmark

We benchmarked the ability of the V-region retrieval service to retrieve highly similar matches with sequence identity 90% or greater. We found 189/738 (25.60%) heavy chains that match greater than 90% sequence identity, with 4/738 (0.54%) perfect matches (Dusigitumab, Edrecolomab, Melredableukin and Zanolimumab). We find 288/707 (40.73%) light chains with sequence identity 90% or greater and 50/707 (7.07%) perfect matches. Therefore, despite restrictive length constraints that produces more relevant results, AbDiver identifies high quality sequence matches.

We benchmarked the ability of our clonotype service to retrieve sequences sharing the same V gene and with CDR3 sequence identity with the query of at least 80% (Greiff *et al*., 2015). For 409/686 (59.62%) therapeutic heavy chains with unique CDRH3s we found matches greater than 80% sequence identity and for 172/686 (25.07%) matches greater than 90%. In 35/686 (5.10%) instances, a combination of V gene and CDRH3 can be matched to an identical CDRH3 and V gene of an NGS sequence. For 384/573 (67.01%) therapeutic light chains with unique CDRL3s we found matches greater than 80% sequence identity and for 279/573 (48.69%) matches greater than 90% with 244/573 (42.58%) perfect matches in V gene and CDRL3 sequence. Therefore, our service succeeds at finding high number of relevant matches for most of the therapeutics in our dataset.

## 3 Discussion

We created an online portal that facilitates the navigation of natural antibody diversity. We envisage particular application of the service to enable drawing parallels between natural and therapeutic antibodies (Krawczyk *et al*., 2019; Petersen *et al*., 2021) for the purpose of engineering to remove Post Translational Modification risks while maintaining favorable biophysical properties. For instance, removing a deamidation motif would require one to introduce one of few standard mutations (e.g., NA, QG) and re-assess the function of the antibody. AbDiver can identify sequence-similar candidates from natural origin, increasing the chances that function and immunogenicity will not be compromised. Beyond facilitating liability removal, AbDiver could excavate sequences with potentially better product profiles than the lead therapeutic. Employing the CDR3 or clonality search can accelerate the discovery of clones that share therapeutic properties of the query, yet provide alternatives with potentially better product profiles. We hope that AbDiver will enable research-supporting applications to facilitate the decision making process in rational design of therapeutics during lead optimization.

