## Supplementary File for "AbDiver – A tool to explore the natural antibody landscape to aid therapeutic design"

**Supplementary materials for AbDiver – A tool to explore the natural antibody land-scape to aid therapeutic design.**

**Section 1. Benchmarking the profile retrieval service.**

We benchmark AbDiver's profiling service ability to offer relevant information for our 742 therapeutics. This took the form of assessing for how many of these, we could not find profiles with sufficient number of sequences (arbitrarily set at 10,000 sequences building the profile) and whether the said therapeutics indeed do not agree with the NGS distribution in their framework regions – thus indicating being engineered-away from the genetic 'centroid'. We plotted the number of sequences contributing to the NGS profiles in Figure 1 for gene-based profiles and Figure 2 for allele-based profiles. We also plotted the number of IMGT framework positions that do not match with top amino acids in identified gene-based (Figure 3) and allele-based (Figure 4) profiles, indicating that for majority of therapeutics, there exist mutations not agreeing with observed natural distribution. The higher number of framework positions disagreeing with NGS distribution in the gene-based profiles reflects multiple alleles contributing to these.

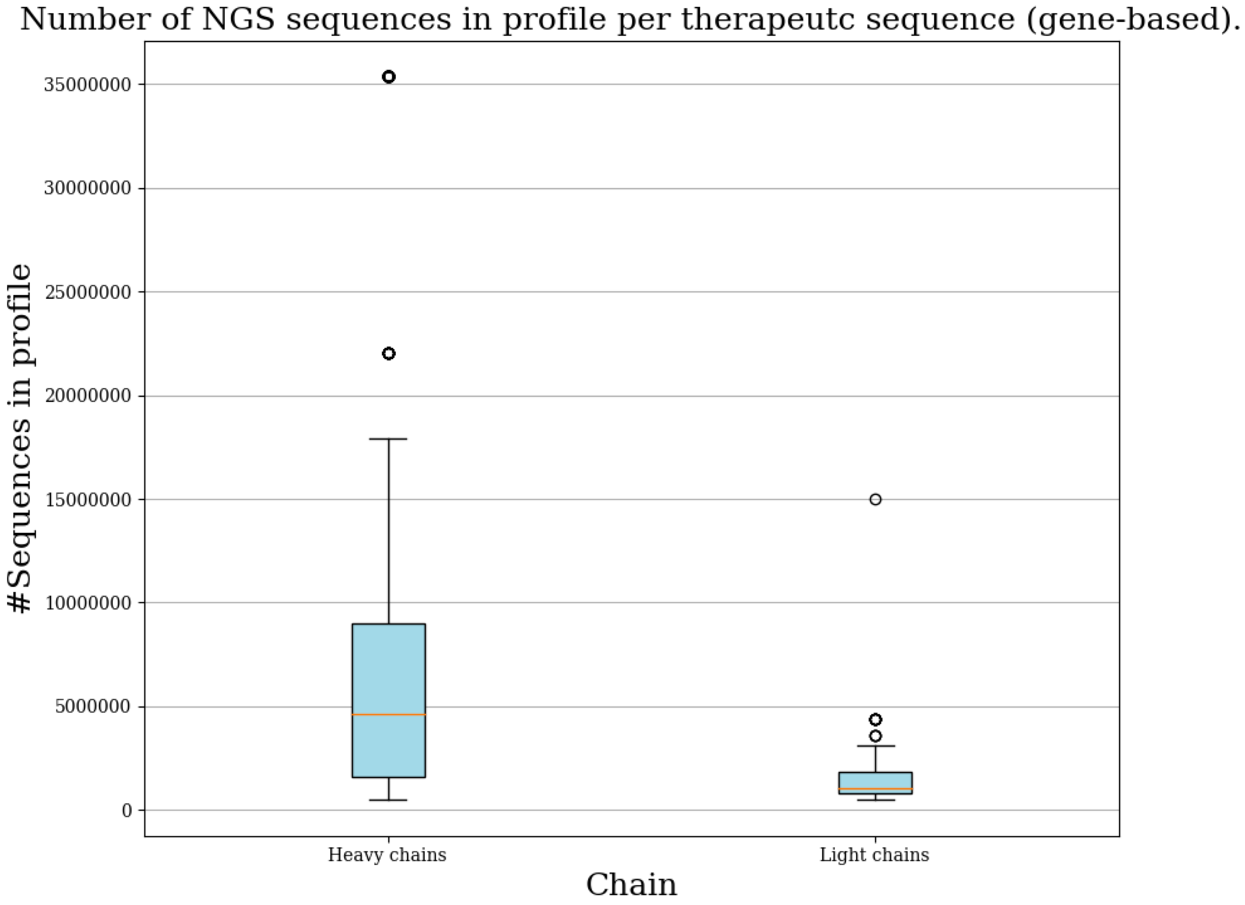


**Figure 1. Number of sequences in gene profiles per therapeutic sequence**. For heavy and light chains in 742 therapeutic antibodies we plotted the number of sequences contributing for a gene-based profile.

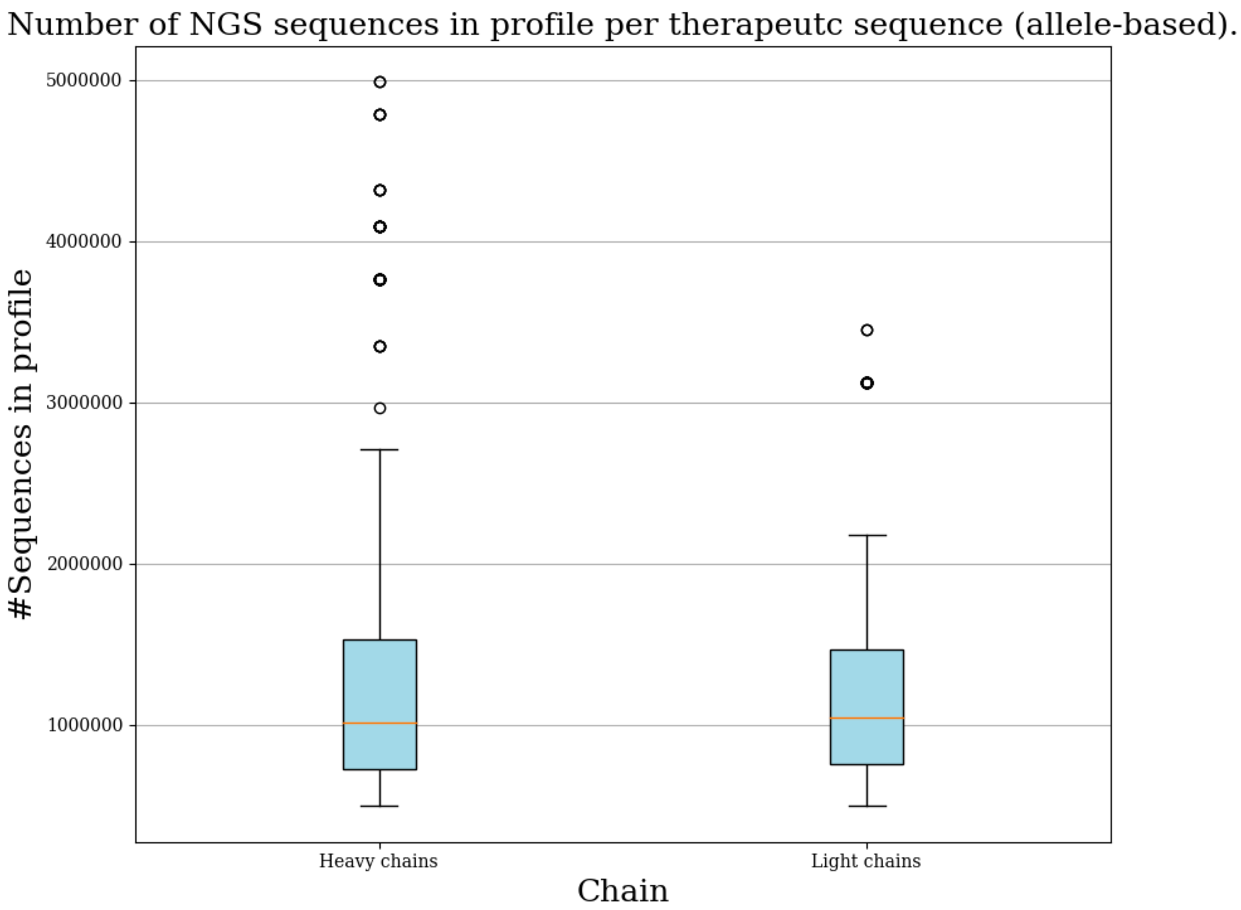


**Figure 2. Number of sequences in allele profiles per therapeutic sequence**. For heavy and light chains in 742 therapeutic antibodies we plotted the number of sequences contributing for a gene-based profile.

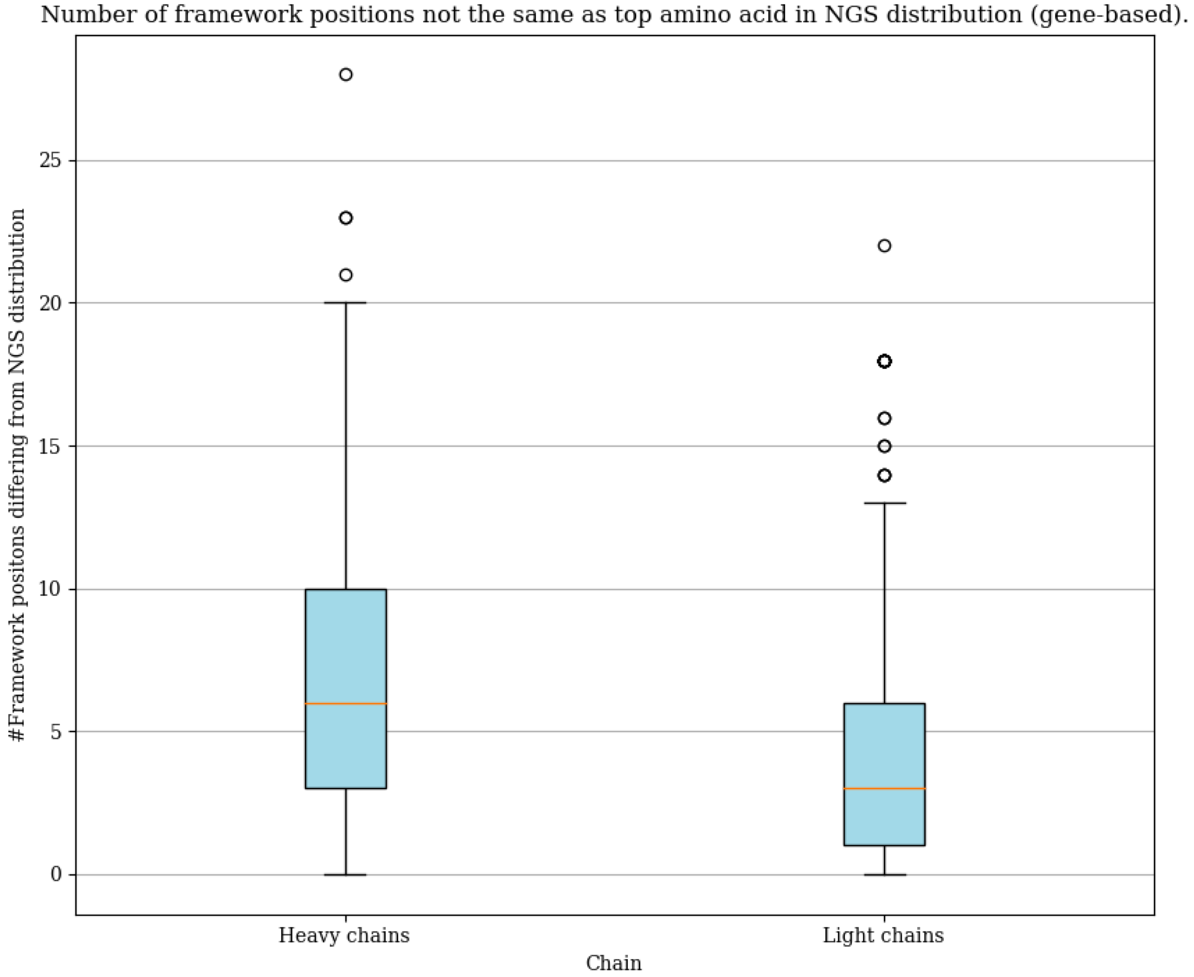


**Figure 3. Number of framework positions not agreeing with top amino acids in gene-based profiles**.

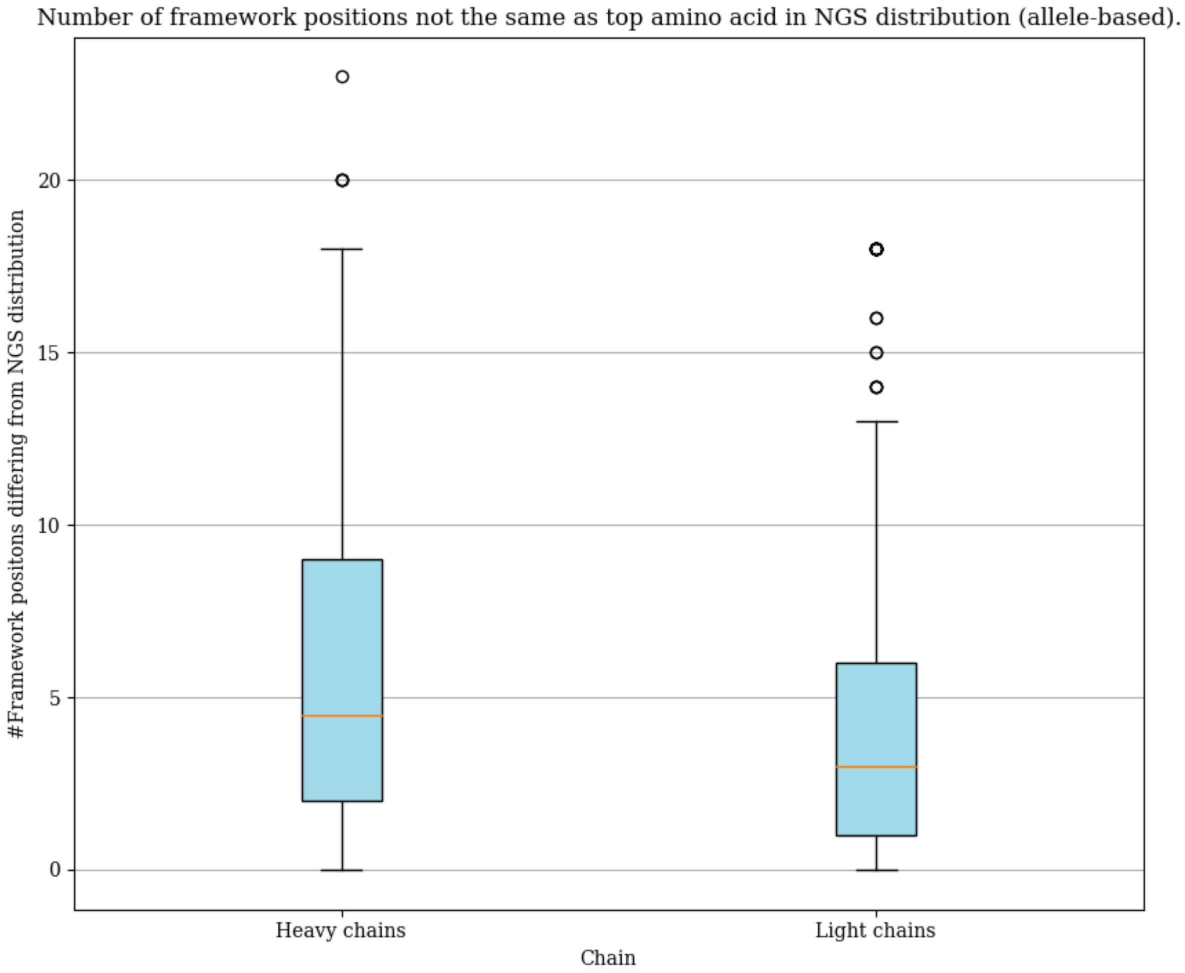


**Figure 4. Number of framework positions not agreeing with top amino acids in allele-based profiles**.

**Section 2. Time benchmark of AbDiver.**

Retrieval time for AbDiver was calculated as time (in seconds) taken from issuing a query to the database, to returning the top 100 results. We benchmarked retrieval of the entire variable region (heavy and light separately) as well as CDR3 retrieval (CDR-H3 and CDR-L3 separately) for 742 therapeutic antibodies. The benchmark, expressed in seconds is given in Figure 5. Mean retrieval time for full sequences was 1.56s for light chains and 4.40ss for heavy chains. For CDR3 retrieval, the average time for CDR-L3 clones was 1.24s and for CDR-H3 clones it was 2.68s.

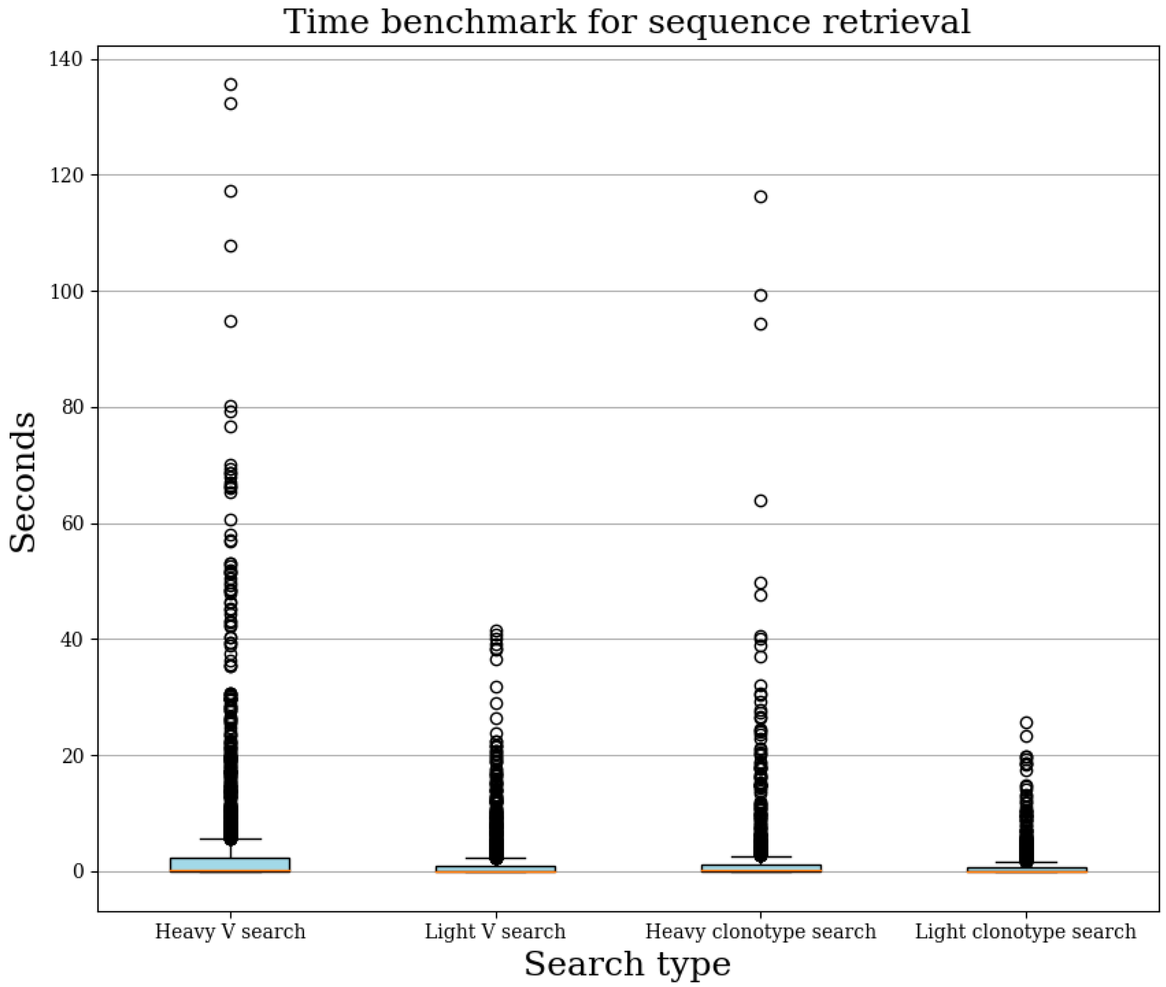


**Figure 5. Retrieval times statistics.** For each chain and search type, the retrieval corresponds to wall clock time from issuing the query to when all the results were processed and returned.
